# OrthoBrowser: Gene Family Analysis and Visualization

**DOI:** 10.1101/2024.08.27.609986

**Authors:** Nolan T. Hartwick, Todd P. Michael

**Author notes:** **Declaration of interest:** T.M. is co-founder of Cquesta, a company that works on crop root growth and carbon sequestration.

## Abstract

The analysis of gene families across diverse species is pivotal in elucidating evolutionary dynamics and functional genomic landscapes. Typical analysis approaches often require significant computational expertise and user time. We introduce OrthoBrowser, a static site generator that will index and serve phylogeny, gene trees, multiple sequence alignments, and novel multiple synteny alignments. This greatly enhances the usability of tools like OrthoFinder by making the detailed results much more visually accessible. This interface can scale reasonably up to hundreds of genomes, allows a user to filter this large dataset to a subset of samples they are interested in at that particular moment in time, or “zoom in” to only a subtree of the orthogroup. The multiple synteny alignment method uses a progressive hierarchical alignment approach in the protein space using orthogroup membership to establish orthology. Orthobrowser makes it easy for users to identify, interact with, explore, and share key information about their gene families of interest.

**Availability:** OrthoBrowser is available under the MIT license at: https://gitlab.com/salk-tm/orthobrowser

**Supplementary Information:** Example OrthoBrowser Results: https://orthobrowserexamples.netlify.app/

## 1. Introduction

Gene family analysis is one of the fundamental tools of modern genomics. By comparing putative protein sequences against each other, we can derive insights including: 1) identification of conserved putatively functional domains; 2) elucidation of potential gene model errors; 3) characterization of orthologous genes, allowing for putative transfer of findings in one genomic context to another not yet studied context; 4) estimation of the phylogenetic history of a group of samples; and 5) exploration of tandem duplication events, deletion events, and whole genome duplication events.

Tools like Orthofinder (Emms and Kelly 2015, 2019) represent state of the art for doing gene family analysis at scale. Orthofinder operates by comparing proteomes for many samples against each other, identifying orthogroups, a set of genes thought to be descendant from some putative ancestor gene, constructing individual trees representing the evolutionary history of each orthogroup, and then estimating an overall phylogeny for the provided samples. Orthofinder produces a variety of useful output files including but very much not limited to: a main orthogroups table, a gene tree for every orthogroup, a phylogeny tree (Emms and Kelly 2019).

Orthofinder output is extremely useful, but requires some degree of expertise to understand, and further computational experience to handle jumping around between files and using a variety of other software to interact with or visualize those results. A typical examination of a gene of interest may require first using grep or some table browsing software like excel to parse through the main orthogroups table and identify the orthogroup containing the gene of interest. The scientist would then need to use some type of tree viewing software like ete3’s (Huerta-Cepas, Serra, and Bork 2016) command line utilities or a larger application like MEGA (Tamura et al. 2013) for loading the tree. Gene trees for non-trivial orthogroups are often large and complicated and frequently require virtual pruning in order to simplify the tree for visualization purposes. If you are working at the command line, this might mean writing a custom script in order to execute the transformation that you need. If you are working in MEGA, it might mean spending several minutes interacting with its robust tree editing tools. At that point it is usually extremely beneficial to view an MSA in order to aid in evaluating the gene tree. There may be duplication events that you want to understand, which usually means using a genome browser and interacting with other file formats like gff3 in order to determine if a duplication is tandem or displaced.

OrthoBrowser seeks to simplify this process through the indexing of a user’s data and the creation of a static website for browsing that data. OrthoBrowser is easy to install through pip and can be run on any Orthofinder result directory, though is not restricted to Orthofinder. It features four main figure types: a sample/species phylogenetic tree, a gene/orthogroup level phylogenetic tree, multiple sequence alignment and multiple synteny alignment. OrthoBrowser is conceptually similar to OrthoVenn3 (Sun et al. 2023), in that it is designed to make it easier for people to interact with phylogeny and gene trees and related information.

OrthoVenn3 fully integrates and manages the underlying analysis, whereas OrthoBrowser is designed to work downstream from tools like Orthofinder. The public OrthoVenn3 instance is limited to 12 samples, and while the local version has no such limit, the interface is clearly designed for working with a handful of samples at a time. OrthoBrowser does not have these limits and scales reasonably well to hundreds of samples. Running OrthoVenn3 locally requires docker, while OrthoBrowser can be easily installed with pip. OrthoBrowser outputs a simple static website making the results more portable, easier to share, and easier to host publicly if desired.

OrthoBrowser makes it easier and faster for scientists to get more information about the genes they actually care about as well as making it easier to share their findings in combination with static web hosting services like github pages or netlify.

## 2. Materials and Methods

OrthoBrowser is implemented in a combination of Python, javascript, html, and css. Essentially, there is a pip installable python package that uses bundled javascript, css, and html templates to process a users input files and produce indexed output files that the bundled html/javascript/css page knows how to search and read. The Python module relies on the etetoolkit (Huerta-Cepas, Serra, and Bork 2016) for processing of newick trees and Biopython (Cock et al. 2009) for fasta format data. The main page relies on a combination of D3 and Plotly (“Data Apps for Production,” n.d.) for generating interactive visualizations alongside bootstrap for other GUI elements.

OrthoBrowser applications consist of two virtual pages. The first displays an overall phylogeny for your samples. The second includes support for three different visualizations: Gene Tree, Amino Acid MSA, and Multiple Synteny Alignment (Figure 1; Supplemental Figure 1). The Phylogeny tree, Gene Trees, and Amino Acid MSA are essentially just visualizations of a user’s provided input. A user must provide a set of gene trees and a phylogeny tree. Amino Acid MSA files are optional but required to generate MSA visualizations. In order to generate the Multiple Synteny Alignment visualizations, users must provide gff3 or bed format files specifying where genes are located in the relevant genomes.

**Figure 1:**
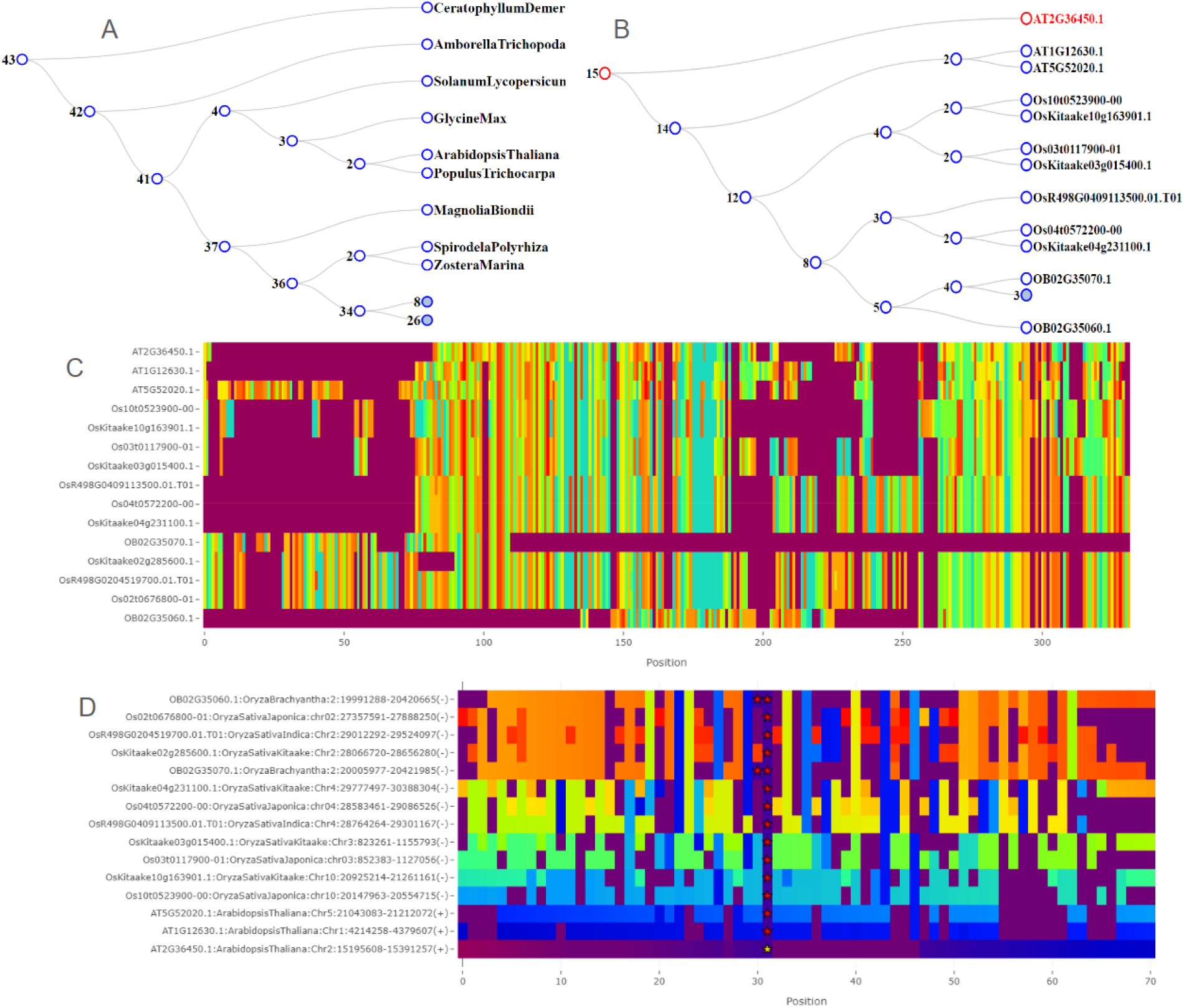
An overview of the figure types available within OrthoBrowser. (A) orthofinder’s estimated phylogeny representing the samples under study: in this case a set of monocot and eudicot genomes collected from Plaza Monocots v5. Trees in OrthoBrowser are interactive and auto-collapse at a preset depth. Interacting with an internal node will zoom in to that subtree. (B), (C), and (D) are figures associated with the *HARDY* (*HRD*) gene highlighting a probable misprediction (split-gene) of the *HRD* in *Oryza brachyantha*. (A) Orthofinder’s estimate of the gene tree associated with *HRD* after subsetting to several rice samples and arabidopsis. The searched gene (*HRD*) is highlighted in red. (C) A visualization of the amino acid (AA) multiple sequence alignment (MSA) associated with the selected genes. Every cell represents an AA or gap. OB02G35070 and OB02G35060 appear to match distinct portions of *HRD* in Arabidopsis and the other rice assemblies. The figure supports zooming and panning and will specify which specific AA or gap is represented by hovering over a cell. (D) A visualization of the local context of the proteins around *HRD* and orthologues. Every cell represents a protein or gap and is colored according to which orthogroup (and orthogroup cluster) the protein belongs to. The searched gene is marked with a gold star. Other genes in the searched genes orthogroup are marked with red stars. Consistent banding is indicative of synteny as observed between Os10t0523900 and OsKitaake10g163901. OB02G35070 and OB02G35060 are adjacent implying a tandem duplication, or more likely a split gene model given the AA MSA. The figure supports zooming and panning and will specify which specific protein or gap and its orthogroup is represented by hovering over a cell.

The Multiple Synteny Alignment uses a hierarchical Needleman–Wunsch alignment approach (Needleman and Wunsch 1970) operating in the protein/orthogroup space rather than the nucleotide or amino acid space. The region around each gene, meaning a user specified number of proteins upstream and downstream from the gene, gets treated as a string of tokens. Two tokens in different strings are said to match if they belong to the same orthogroup. They have a perfect match if they belong to the same orthogroup sub-cluster. Two proteins belong to the same orthogroup sub-cluster if they are in the same orthogroup and there exists an internal node that is an ancestor of both proteins less than a user specified total edge length apart. When performing the hierarchical alignment and comparing two positions in two MSA, the sub positions are compared in a pairwise manner and the total score is summed. The Gene tree is repurposed here to act as a guide tree for the hierarchical alignment.

In addition to the main four visualizations, OrthoBrowser applications include several key interface features: 1) a robust search feature with auto-complete allowing users to search for any protein of interest in the dataset; 2) a dropdown menu allowing for selecting a subset of samples of interest, which will filter the relevant trees and MSA dynamically; 3) two buttons allowing for exporting the currently selected protein set in fasta or the current amino acid MSA; 4) a home button linking back to the main phylogeny; 5) all trees “collapse” after a certain depth ensuring the application stays useful even for large trees. Internal nodes are annotated with a count of leaf nodes associated with the subtree; 6) all trees are interactive. Clicking on an internal node will “zoom in” to the subtree rooted at that node. The path from the tree’s root to the user’s searched gene gets highlighted in red; 7) all MSA visualizations are interactive for zooming/panning and provide more information on hover and can be exported to PNG; and 8) collapsible/expandable captions that provide basic explanations for the figures.

## Conclusion

OrthoBrowser acts as an augmentation of Orthofinder, and similar tools, by providing an integrated and user-friendly platform for gene family analysis and visualization. By simplifying the access to complex datasets and visualizations, OrthoBrowser facilitates deeper insights into evolutionary biology and functional genomics. This tool not only reduces the need for specialized computational skills but also accelerates the process of scientific discovery. Researchers can now quickly identify key features and relationships within gene families, enhancing their ability to share and discuss findings with the broader scientific community.

Future updates and community contributions to OrthoBrowser could further enhance its capabilities and adaptability, making it an indispensable resource in genomic research. In particular, we plan to expand the home page to include summary stats such as “percent of genes in orthogroups” allowing users to more easily evaluate their overall analysis. OrthoBrowser demonstrates the potential of bioinformatics to democratize data analysis, allowing more scientists to contribute to and benefit from the rapidly expanding genomic knowledge base.

## Acknowledgments

This work was supported through the Salk Harnessing Plants Initiative (HPI) with funding from the TED Audacious, Bezos Earth Fund, and Hess Corporation. This research was funded by a Bill & Melinda Gates Foundation (BMGF) grant (INV-040541) to T.P.M., and the Tang Genomics Fund.

