## Supplemental Figure 1 for "OrthoBrowser: Gene Family Analysis and Visualization"

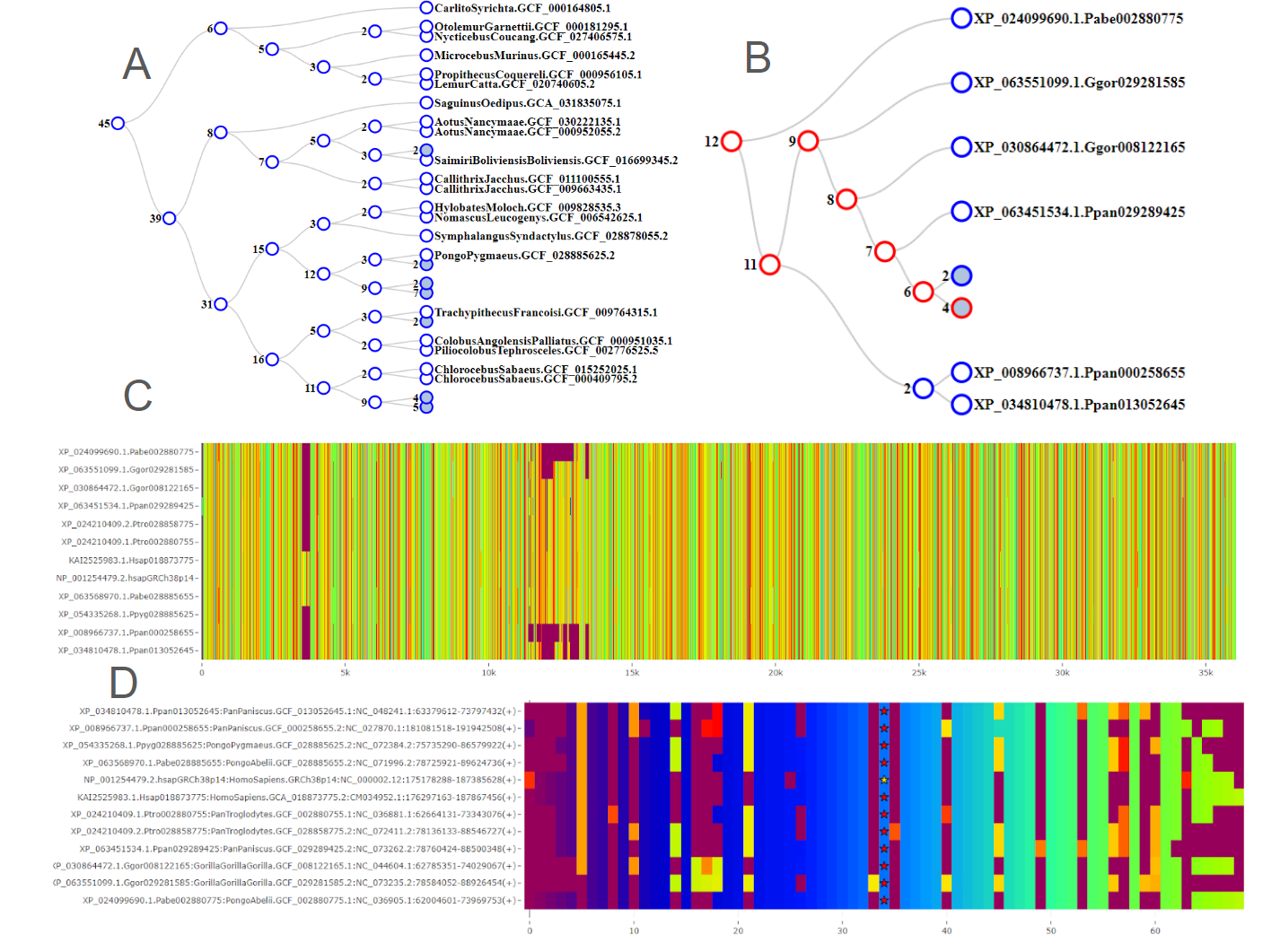
**Supplementary Figure 1: An overview of the figure types available within OrthoBrowser using Primapes**. (A) Orthofinder’s estimated phylogeny representing the samples under study: in this case a set of mammal genomes collected from UCSC Primates (Haeussler et al. 2019). Trees in OrthoBrowser are interactive and auto-collapse at a preset depth. Interacting with an internal node will zoom in to that subtree. (B), (C), and (D) are figures associated with the *Titin* (*TTN*) gene highlighting the OrthoBrowser’s ability to scale to even the largest known proteins. (A) Orthofinder’s estimate of the gene tree associated with *TTN* after subsetting to Hominidae. The searched gene (*TTN*) is highlighted in red. (C) A visualization of the amino acid (AA) multiple sequence alignment (MSA) associated with the selected genes. Every cell represents an AA or gap. The figure supports zooming and panning and will specify which specific AA or gap is represented by hovering over a cell. (D) A visualization of the local context of the proteins around *TTN* and orthologues. Every cell represents a protein or gap and is colored according to which orthogroup (and orthogroup cluster) the protein belongs to. The searched gene is marked with a gold star. Other genes in the searched genes orthogroup are marked with red stars. Consistent banding is indicative of synteny. All proteins under display are syntenic. The figure supports zooming and panning and will specify which specific protein or gap and its orthogroup is represented by hovering over a cell. oment in time, or “zoom in” to only a subtree of the orthogroup. The multiple synteny alignment method uses a progressive hierarchical alignment approach in the protein space using orthogroup membership to establish orthology. Orthobrowser makes it easy for users to identify, interact with, explore, and share key information about their gene families of interest.
